# Discovery of a new fibronectin-binding surface protein of *Streptococcus canis* with serum opacification activity through Transposon Directed Insertion-Site Sequencing

**DOI:** 10.64898/2025.12.05.692515

**Authors:** Miriam Katsburg, Anna Kopenhagen, Etienne Aubry, Mathias Müsken, Gloria Riebesell, Deborah Simmert, Sanja Haake, Inga Eichhorn, Silver A. Wolf, Simone Bergmann, Marcus Fulde

**Affiliations:** Institute of Microbiology and Epizootics, Centre for Infection Medicine, Freie Universität Berlin, Berlin, Germany; Veterinary Centre for Resistance Research (TZR), Freie Universität Berlin, Berlin, Germany; Institute of Microbiology, Technische Universität Braunschweig, Braunschweig, Germany; Central Facility for Microscopy, Helmholtz Centre for Infection Research, Braunschweig, Germany; Genome Competence Centre (MF1), Robert Koch Institute (RKI), Berlin, Germany

**Keywords:** TraDIS, high-throughput screening, fibronectin, serum opacification, endothelial infection, endothelial wound healing, streptococcal adhesin, infective endocarditis

## Abstract

Infective endocarditis is a rare but severe disease in humans and dogs which can be caused by *Streptococcus canis*. The bacterial factors that mediate endothelial adherence remain poorly defined. To understand how *S. canis* can adhere to and invade the endocardium, we combined a high-throughput approach called transposon directed insertion-site sequencing (TraDIS) with a physiologically relevant endothelial infection model that incorporates venous-range shear stress to identify *S. canis* genes required for host cell interaction. A saturated transposon library of clinical strain IMT49926 was screened in a microfluidic infection assay, enabling genome-wide selection of mutants impaired in endothelial adhesion. Comparative analysis of input and non-adherent output pools revealed several candidate genes, including a fibronectin-binding LPXTG-anchored surface protein with high homology to streptococcal serum opacity factors (SOFs). Our findings identified ScSOF as a multifunctional surface protein that plays an important role in the infection potential of *S. canis*. It facilitates adhesion to endothelial cells, prevents endothelial wound closure, contributes to the streptococcal surface architecture, binds fibronectin, opacifies serum, and inhibits β-haemolytic activity. ScSOF represents a strong candidate for future studies into pathogenesis, immune evasion, and (potentially) vaccine or therapeutic targeting in *S. canis* infections.

**Author’s summary:** *Streptococcus canis* is a bacterium commonly found in healthy dogs and cats, but it can sometimes cause serious infections in animals and humans, including infective endocarditis, a dangerous infection of the heart lining. To cause this disease, the bacteria must first attach to and invade the cells that line blood vessels. However, very little is known about how *S. canis* can do this. In this study, we used a genetic screening method that includes all the genes from *S. canis* to identify which bacterial genes are needed for attachment to human endothelial cells under conditions that mimic real blood flow. We discovered a previously uncharacterized surface protein, which we named ScSOF, that plays several important roles during infection. ScSOF helps the bacteria bind to fibronectin, a major host tissue protein, alters the bacterial surface structure, and causes serum opacification, a known marker of virulence in related streptococci. When ScSOF was removed, the bacteria were much less able to attach to endothelial cells, cause cell damage, or interfere with the healing of endothelial wounds. Our findings show that ScSOF is a key factor that enables S. canis to interact with host tissues and may contribute to heart valve infections.

## Introduction

*Streptococcus canis* (*S. canis*) is an opportunistic pathogen that inhabits mucosal surfaces in up to 6.5% of healthy dogs but has been implicated in as many as 22.4% of streptococcal infections in dogs and even up to a quarter of canine infective endocarditis cases [1–3]. *S. canis* has been shown to be the cause of several cases of infective endocarditis (IE) in human patients as well, often after infection of a wound due to close contact with pet cats or dogs [4–8]. IE is a life-threatening condition in both human and veterinary medicine, marked by a high mortality rate especially in canines due to difficulty in diagnosis and treatment [8–10]. Typical IE pathogenesis starts with entry of the bacteria into the bloodstream via wounds or surgical intervention sites [11]. Some of the bacteria may escape phagocytosis in the bloodstream and can adhere to the endocardium to establish infection. Adherence to the endocardium can be established through direct interactions with the host endothelial cells or interactions with the host matrix proteins. The endocardium displays an increased vulnerability to infection due to prior damage in the endothelial layer, caused by common risk factors for IE such as obesity and age. In addition, mechanical stress mediated by high blood pressure is often responsible for endothelial damage. Other causes are inflammation and infection [12].

Once the bacteria colonize the endocardium, they can form a complex biofilm, which is typical for IE and results in treatment difficulties. According to Lerche et al., disruption of the biofilm could improve the treatment outcomes of IE cases [13].

One contributing factor to adhesion and invasion of the host cells is the *S. canis* M protein SCM, which was previously implicated in IgG binding and immune evasion and thought to be involved in early colonization stages through interactions with host factors such as fibrinogen and plasminogen, depending on the M protein type [1416]. However, the contribution of SCM to virulence in an in vivo murine model and an in vitro vaginal epithelial cell adherence assay was modest [17]. The molecular mechanisms by which *S. canis* adheres to host tissue and initiate biofilm formation remain largely uncharacterized.

In *Staphylococcus aureus*, the pathophysiology of infective endocarditis has been described in greater detail. Here, it is known that the MSCRAMMs, microbial surface component reacting with adhesive matrix molecules, have a crucial impact in the ability of bacteria to interact with host matrix proteins and cause endocarditis.

Important functions of these factors are fibronectin-binding, fibrinogen-binding and platelet-binding [18]. In another study it was also shown that fibrinogen-binding proteins of *S. aureus* work together with fibronectin-binding proteins to enable the colonization of heart valve tissue and the invasion of the endothelium *in vivo* [19]. A similar role of fibronectin-binding proteins was also described in streptococci, with *Streptococcus mutans* as a common causative bacterium in human IE and *Streptococcus pyogenes* as an uncommon causative pathogen [20, 21]. Moreover, it was described that fibrinogen bound to the surface of *S. mutans* increased adherence to endothelial cells and could act as a bridging molecule to mediate biofilm formation [22].

To elucidate the mechanism of adherence to the endothelial cells, a crucial step in the development of IE, we aim to identify key bacterial adherence factors of *S. canis* by using transposon directed insertion-site sequencing (TraDIS) as a high-throughput sequencing approach. TraDIS has already been successfully applied to identify essential bacterial genes in various growth conditions, such as oxidative stress or antibiotic pressure [23–26]. Previously, this technique was also used to identify virulence genes in various *in vivo* infection models [27, 28]. In accordance with the 3R principle, animal experiments should be replaced as much as possible by *in vitro* assays. Modern model systems allow us to grow cells in conditions that are comparable to the living host. This study is the first report of using TraDIS in combination with an *in vitro* endothelial cell infection assay under simulation of physiological blood flow to study relevant gene products for adherence and invasion.

We established an *in vitro* infection model that closely reflects the dynamic blood flow environment encountered by endothelial cells within the vasculature. Endothelial cells are constantly influenced by hemodynamic forces, particularly shear stress generated by blood flow, which governs their structural and functional characteristics. To replicate these physiological conditions, we employed a microfluidic system (Ibidi®) capable of applying defined shear stresses (6–10 dyn/cm²), thereby simulating the venous flow environment relevant for bloodstream infections such as endocarditis and septicaemia. This setup allows us to directly compare bacterial adherence and host cell responses under both flow and static conditions, enabling the identification of adherence factors that are specifically required for colonization under physiologically relevant shear stress. The use of microfluidics ensures consistent and reproducible flow conditions, supporting the formation of endothelial monolayers that closely mimic *in vivo* behaviour and providing a robust platform for functional genomics screens such as TraDIS [29].

In this study, we identified a surface protein of *S. canis* called ScSOF, the *S. canis* serum opacification factor. We constructed an isogenic knock-out mutant and compared the ability of this mutant to infect endothelial cells under static and flow conditions with the wildtype strain. We microscopically visualized the morphology and surface structure of the mutant using electron microscopy and tested the ability of the mutant and the wildtype strain to bind to human fibronectin and opacify serum. We also investigated the influence of infection with the wildtype or mutant strain on endothelial wound healing.

## Materials and Methods

### Bacterial strains and growth conditions

In this study, *S. canis* strain IMT49926 and its isogenic mutant were used. Strain IMT49926 was isolated from a case of canine infective endocarditis during routine diagnostics at Freie Universität Berlin [9]. For serum opacification activity analysis, *S. suis* strain 10 was used as a positive control [30]. Streptococci were grown on Columbia agar plates with 5% sheep blood (BD) or in Todd Hewitt broth (THB) (Roth) unless indicated otherwise. *Escherichia coli* strains were cultured in Luria-Bertani (LB) medium. In appropriate cases, antibiotics were added at the following concentrations: erythromycin (Erm), 150 µg/mL for *E. coli*, 2 µg/mL for *S. canis*.

### Generation of transposon library

*S. canis* IMT49926 was transformed with plasmid pGh9:IS*S1* by electroporation and a transposon library was generated as described previously [26].

### *In vitro* HUVEC infection assays

Primary HUVECs (PromoCell, Germany) were cultured in endothelial cell growth medium (C-22010, PromoCell, Germany) and infected with mid-log phase *S. canis* TraDIS library, *S. canis* WT or ScSOF knockout strains at multiplicity of infection (MOI) 10. For flow conditions, the Ibidi µ-Slide I Luer system was connected to a software-controlled pneumatic pump (Ibidi, Germany) to simulate physiological shear stress conditions on endothelial cell surfaces. Cells were grown to confluency under flow conditions of 10 dyn/cm² and then infected for 2.5 hours at shear rates of 6 dyn/cm². Following infection, cells were washed, fixed using 3% paraformaldehyde in PBS, and stained to assess both, bacterial association and cytoskeletal integrity of the host cells. The cytoskeleton was visualized using Alexa Fluor 488-conjugated phalloidin, while cell nuclei were counterstained with DAPI. *S. canis* bacteria were detected using a *S. canis-*specific primary antibody. Overlays of different fluorophore signals were used to distinguish between attached and internalized bacteria by first using an Alexa Fluor 488 secondary antibody to stain only extracellular bacteria, followed by an Alexa Fluor 568 secondary antibody after permeabilization with 0.01 % Triton X-100. Infection analyses with the *S. canis* TraDIS-library were repeated twice with the aim to increase the selective pressure, thereby enforcing a sharp discrimination between those bacteria of the library, which were tightly associated with the host cells, from those, which lost the ability to attach to the host cells due to transposon integrations. The repetition of cell infection analyses resulted in bacterial output pools named the first and the second selection.

In static infection assays, HUVECs were grown to confluency on 12 mm glass coverslips in a 24-well plate. Cells were then infected at various MOIs ranging from 0.5 to 5 for 1 hour, followed by replacement of the medium and incubation for an additional 2 hours. Immunofluorescent staining was performed as previously described. Images were acquired as z-stacks using a confocal laser scanning microscope (CLSM; Leica SP8) equipped with LAS X software (Leica, Germany), employing a 63x/0.75 oil immersion objective lens to achieve a total magnification of 630x.

### *In vitro* A549 infection assays

A549 lung epithelial cells were cultured in DMEM supplemented with 10% FBS. Cells were cultured at 37°C and 5% CO_2_. Cells were seeded (1*10^4^ cells/well) on glass coverslips in a 24-well plate, grown to confluency and subsequently infected with *S. canis* WT or ScSOF knockout strains at MOI 5 for 2.5 hours. Post-infection, cells were washed, fixed, and stained for bacterial association and host cytoskeletal integrity using rabbit anti-*S. canis* polyclonal antibody, goat anti-rabbit Alexa Fluor 488 and Alexa Fluor 647 after permeabilization as well as Alexa Fluor 580conjugated phalloidin as described in the HUVEC infection assay. DAPI was used to stain cell nuclei. For quantification, a similar infection was executed without the glass coverslips. Bacteria-infected wells were washed twice with PBS and lysed by 1% (w/v) saponin in H_2_O. Serial dilutions in PBS were plated out on THB-Agar. Colonies were counted after incubation for 48 hours at 37°C.

### Chamber-separation cell migration assay (CSMA)

Endothelial gap closure was assessed using the CSMA as described previously [31]. In short, HUVECs (3*10⁵ cells/mL) were seeded in Ibidi® three-chamber silicone inlets on gelatine-coated dishes and grown to confluence. After seeding and reaching confluence in Ibidi® inlets, cells were exposed to increasing shear stress (3–10 dyn/cm²) and maintained at 10 dyn/cm² for up to 48 h. After inlet removal, a standardized 500 μm gap was created. Gap closure and cell morphology were monitored microscopically at multiple time points, with actin and nuclei visualized by DAPI and Alexa Fluor 488-conjugated phalloidin. Images were obtained by CLSM (SP8, Leica) and cell-free areas were quantified and normalized to baseline using LAS X software. To study the effects of *S. canis* infection, HUVECs were preconditioned under flow, followed by infection with *S. canis* or its ΔScSOF mutant (MOI 5) under continuous shear stress. Media were exchanged periodically to limit bacterial overgrowth. Experiments were performed in triplicate and analyzed for effects on endothelial migration and morphology.

### Differentiation between proliferating and migrating cells during gap closure

EC proliferation and migration during gap closure were assessed using the EdU click assay in the CSMA setup, with 10 μM EdU labelling proliferating cells [31]. After fixation, EdU-positive (red) and migrating (blue) nuclei were visualized by fluorescence and quantified by CLSM (SP8, Leica). Nuclei were counted in randomly selected areas within the 500 μm gap at multiple time points. All experiments were performed in triplicate, and results are reported as mean ± SD.

### DNA preparation and sequencing

DNA was extracted from the mutant libraries after overnight growth at 37 °C in Todd Hewitt Broth. DNA was quantified using the Qubit dsDNA HS Assay and 1 µg of input DNA was used to start the library preparation. Fragmentation, A-tailing, and end repair, was performed using the NEBNext Ultra II FS DNA Library Prep Kit for Illumina (NEB #E7805, E6177), yielding fragments of approximately 400 bp.

A Y-adaptor was generated in-house using Illumina multiplexing adaptor sequences (Illumina) according to previous methods [26]. Adaptor ligation was executed by adding 15 µM of the adaptor to the fragmentation mixture, followed by NEBNext Ultra II Ligation Master Mix and NEBNext Ligation Enhancer. Samples were incubated at 20°C for 15 minutes in a thermocycler.

Purification by AMPure Xpbeads, PCR amplification, PCR clean up, and size selection were done following the method described by Charbonnaeu et al [26]. Final library concentration was measured using the Qubit dsDNA HS Assay and adjusted to yield 10 pM final load concentration for the pooled replicates.

To prepare for sequencing, the NextSeq™ 2000 P3 Reagents (200 Cycles) were used from Illumina. The libraries were combined with 40% PhiX (Illumina). Sequencing primers were used as described by Charbonnaeu et al [26]. The amplified libraries were single end sequenced using the Illumina NextSeq at the Genome Competence Centre of the Robert Koch Institute.

### TraDIS analysis

For TraDIS analyses, genome sequences from the *S. canis* library used for infection analyses are named “input pool” and sequences from bacteria recovered after cell culture infections are named “output pool”. From input pool and from all output pools, raw fastq files were analysed using the Bio-TraDIS (v1.4.5) scripts of the Sanger Welcome Trust Institute (https://github.com/sanger-pathogens/Bio-Tradis) in a similar way as Charbonneau et al [25, 26]. First, contigs were trimmed to 68 bp using the Trimmomatic read trimming tool (v0.40) [32]. Next, the bacteria_tradis pipeline script filtered reads according to the specified transposon tag (GAGAAAACTTTGCAACAGAACC). After tag removal, the remaining 46 bp of *S. canis* DNA were mapped to the IMT49926 reference genome (NCBI Bioproject: PRJNA945807) using SMALT short read mapper (v0.7.6) (http://www.sanger.ac.uk/resources/software/smalt/), producing a plot file of insertion sites for downstream analysis. Insertion sites were plotted on the reference genome using DNAplotter in the Artemis genome browser (v18.2.0) [33, 34]. A transposon tag mismatch of 1 was allowed, while the mapping threshold was set to 100%. Next, the command tradis_gene_insert_sites was used to create a document of unique insertion sites, total read counts and insertion indices. The output file from tradis_gene_insert_sites was used in tradis_essentiality to determine the essential genome of *S. canis* strain IMT49926. The essential genes were later excluded from the list of potential adhesion factors. Lastly, the tradis_comparison.R script was used to compare the input (control) and output pools (conditions) using the tradis_gene_insert_sites output files from each experimental replicate (n=3) of the pools. This generates a volcano plot illustrating log 2-fold change and -log2 (qvalues) for every mutagenized gene as well as a csv file summarizing this information. The csv files were used for further downstream analysis in R.

The IMT49926 genome was functionally annotated using the eggNOG-mapper 2.1.12 [35]. COG annotations were subsequently added to the csv output files from the TraDIS analysis. To visualize the data, the csv file was analysed with R (v4.4.0). A subset of hits made of the genes with logFC>2 and q value<0.01 was selected. This subset was used to create a heatmap with pheatmap. A second subset was designed to compile only genes with an LPXTG-anchoring domain, which are characterizing surface proteins of Gram-positive bacteria. Analogously, a heatmap was created from this subset.

### Targeted mutagenesis of ScSOF

Targeted mutagenesis of ScSOF was performed as described previously but using a newly constructed pGh9:ΔSOF vector [16]. The regions 500 bp upstream and downstream of the *sof* gene were amplified from gDNA of IMT49926 using the primer pairs: 5’-GGCCATGGTAGGGGAACTAGAC-3’; 5’-CCCTATAGCCGACAGATAATTTCC-3’ and GGGATATCGGAGCAGCAGGAC; 5’GGCCATGGGACGTTCATTTTGAG-3’. The two PCR products were digested using EcoRV and ligated with T4 Ligase. The resulting 1 kb region was amplified again with the previous primers 5’-GGCCATGGTAGGGGAACTAGAC-3’ and 5’-GGCCATGGGACGTTCATTTTGAG-3’. The product was digested with NcoI and inserted into the previously described pGh9:ΔIS*S1* vector [16]. 200 ng of the resulting plasmid was introduced into IMT49926 by electroporation. Selection and verification of the mutant was performed as described prior. The isogenic mutant was confirmed using PCR using the primer pair mentioned above and through WGS.

### Electron microscopy

For imaging of liquid bacterial samples, *S. canis* IMT49926 and 49926Δ*sof* were grown to an optical density (OD_600nm_) of 0.4 before being fixed. The fixative solution contained (per sample): 2880 μL cell culture medium, 800 μL paraformaldehyde (25%), and 320 μL glutaraldehyde (25%). Sample preparation and imaging for scanning electron microscopy (SEM) and transmission electron microscopy TEM were performed at the Central Facility for Microscopy (ZEIM) at the Helmholtz Centre for Infection Research and performed as described before [31, 36].

### Fibronectin binding by FACS

*S. canis* IMT49926 and 49926Δ*sof* were grown to mid-exponential phase in TSB medium. Cells were washed once in PBS and adjusted to 1*10^8^ CFU/mL. A total of 2*10^7^ CFU was suspended in 100 µL PBS with 0.5% BSA. Fibronectin-FITC or Fibrinogen-FITC (positive control) was added to obtain a final concentration of 50 µg/mL. Bacteria were incubated in the dark at 37°C for 30 minutes. Bacteria were then washed in PBS and fixed in 4% paraformaldehyde in PBS. A total of 10^6^ bacteria were analysed by flow cytometry as described before [16].

### Fibronectin binding on glass coverslips

In a 24-well plate, 12 mm glass coverslips were inserted and coated with 50 µg/mL of human recombinant fibronectin. After washing with PBS, a bacterial culture of either *S. canis* IMT49926 or 49926Δ*sof* with OD 0.05 in THB was added to the wells. After the indicated incubation time, unbound bacteria were aspirated and wells were washed at least three times with PBS, after which they were fixed with 4% paraformaldehyde in PBS for 30 minutes. Bacteria were stained with polyclonal anti-*S. canis* and secondary Alexa fluor 488 antibodies. After embedding, images were obtained using a DMI6000B fluorescence microscope equipped with a 63x/1.40 oil immersion objective and operated with LAS X software (Leica, Germany). The surface area covered by bacteria was quantified by ImageJ.

### Serum opacification activity

For excreted SOF and SOF that is covalently linked to the cell wall, *S. canis* IMT49926 and 49926Δ*sof* were grown to mid-exponential phase in 10 mL THB and centrifuged (5000 *xg*, 5 min) to separate supernatant from the cell pellet. The cell pellet was resuspended in 1 mL of sterile PBS. Both samples, supernatant and cell suspension, were heat-treated at 65°C for 20 min to stop bacterial growth. Afterwards, 10 µL sample were added to 100 µL of horse serum in a 96-well plate format. After overnight incubation at 37°C, 100 µL of 0.85% sterile NaCl-solution was added and absorbance measured at 450 nm. For non-covalently associated SOF, 1% SDS extracts of overnight bacterial cultures were prepared as described by Courtney and Pownall [37]. 25 µL of SDS extract was added to 100 µL of horse serum in a 96-well plate and incubated at 37°C overnight before absorbance was measured as mentioned above. Strain *S. suis* M10 was used as a positive control, while horse serum alone acted as a negative control.

### Statistical analysis

Graphs and statistical calculations were generated using the Prism software (v8.0) (GraphPad). Unpaired, two-tailed student t tests were performed unless specified otherwise in the figure captions. * Indicates p < 0.05, ** indicates *p* < 0.01, *** indicates *p* < 0.001 and **** indicates p < 0.0001.

## Results

### Identification of novel *S. canis* IMT49926 genes involved in adherence and invasion of endothelial cells

The clinical *S. canis* strain IMT49926 was isolated from a case of infective endocarditis [9] and was used for the generation of a genomic transposon library displaying 26056 unique insertion sites. Library saturation and insertion density is plotted in Fig 1A, showing genome wide coverage of the transposon insertions on average every 75 bp. Using Artemis for visualization, transposon insertion into every gene was manually confirmed. The library was used to infect a for cell culture under defined flow conditions and thus, identify transposon mutants with reduced cell adherence ability. For this purpose, two rounds of infection were subsequently performed providing a first and a second set of separately harvested adherent and non-adherent bacteria as output pools. Confirmation of the successful separation of adherent and non-adherent transposon mutants is provided by immunofluorescence microscopy after infection with the input library (first set) and with the non-adherent output pool (second set) as shown in Fig 1B. The immunofluorescence overlay visualizes a reduced number of adherent bacteria after the 2^nd^ round of infection compared to the first (Fig. 1B, yellow arrows). Bioinformatic analyses revealed no sufficient separation of adherent and non-adherent transposon mutants in the output pools after the first infection cycle as most genes had a log 2-fold change (logFC) between -2 and 2 (Fig S1). After the second infection of HUVEC cells with nonadherent bacteria, collected in the supernatant of the first infection round, the separation between non-adherent transposon mutants and its input pool was much clearer, as seen in the heatmap in Fig 1C. Here, bioinformatic analyses revealed that numerous genes showed a significant logFC of over 2 or under -2 in the output pool as clearly seen in the volcano plot in Fig 1D which displays the comparison of read counts of transposon insertions in the non-adherent bacteria harvested after the second round of infection and the input library. To find potential candidate genes for bacterial adhesion, we consider genes that have a significantly positive logFC, meaning that these transposon mutants were found more in the non-adherent output pool. This indicates a disadvantage in adherence or invasion of the endothelial cells for these mutants.

**Fig 1:**
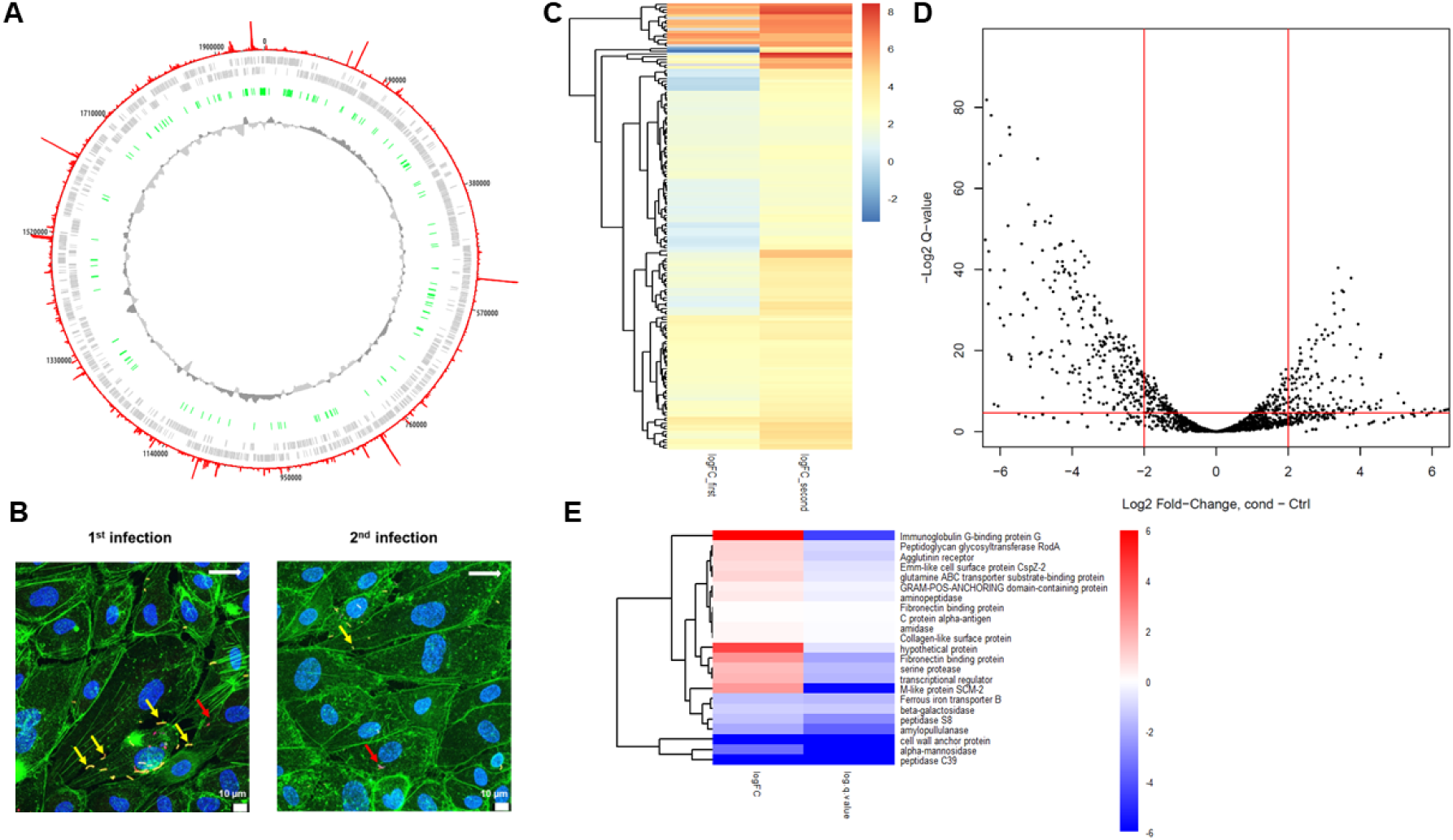
Identification of *S. canis* genes involved in adherence and invasion of endothelial cells. A) Circular genome representation of IMT49926. Transposon insertion frequency is represented in red, gene annotations in grey. In green, the essential genes are depicted, while the inner track illustrates the GC content. Image was created with DNAplotter. B) Representative CLSM image HUVECs grown under flow conditions and infected with the input pool (1^st^ infection) and with the output pool of the first infection round, showing a decrease in adherence (2^nd^ infection). Adherent bacteria are highlighted in yellow and are indicated by yellow arrows. Internalized bacteria are indicated in red/red arrows. HUVEC actin cytoskeleton is marked with phalloidin-Alexa 488 in green and cell nuclei are stained with DAPI in blue; the white arrow at the upper right marks the direction of flow, scale bar indicates 10 µm. C) Heatmap of all significantly overrepresented, mutagenized genes in the output pool after the second selection (right column) with according logFC of these genes after the first round of infection in the left column. Only genes with a q value < 0.01 are considered to minimize the probability of false positives. Genes are clustered according to their profiles in all samples (dendrogram, left side). D) Volcano plot of a comparative analysis of the generated output pool (non-adherent bacteria after second selection round) and the input library showing the logFC of all genes after two rounds of infection compared to the input pool (control). E) Heatmap of all LPXTG-anchoring domain containing genes with the logFC in the left column and log 2 q-value in the right column.

TraDIS-Sequencing of the output pool from the second selection round was followed by selection based on bioinformatic sequence characterization (COGs) and annotation, specifically selecting for genes with an LPxTG-anchoring domain, which is a hallmark of cell-wall anchored surface proteins of Gram-positive cocci [38]. In sum, 23 genes were identified carrying the classical coding sequence for an LPxTG-mediated cell wall anchorage of which 3 genes remained with the criteria (logFC > 2 and q < 0.01) likely encoding proteins relevant for the endothelial infection under theses experimental settings. Based on functional domain homologies and on gene annotations, these 3 genes are identified as an IgG-binding protein G homologue, an annotated fibronectin binding protein, and the M-like protein SCM type II. Since fibronectin-binding proteins are indicated as virulence factors in IE pathogenicity, we selected this gene for further functional investigation [19, 20]. By comparing the gene using the NCBI BLAST core_nt database, this gene showed high similarities (>90%) with serum opacity factors of other streptococcal species. We will refer to the gene in the rest of this manuscript as *sof* and the corresponding protein as ScSOF, the serum opacity factor of *S. canis*. To test the functional characteristics of ScSOF, an isogenic deletion mutant of the *sof* gene was constructed by targeted mutagenesis.

### ScSOF is a serum opacity factor that is covalently and noncovalently associated with the cell surface and plays a role in the inhibition of ß-haemolysis

BLAST analysis showed high protein sequence similarities (>90%) of ScSOF with the LPxTG-motif carrying serum opacity factors of *S. pyogenes* and *Streptococcus dysgalactiae [37, 39–43].* Therefore, we tested the ability of the ScSOF to opacify serum. No serum opacification activity was detected upon incubation of horse serum with bacterial supernatant, indicating that no ScSOF protein is present in the culture supernatant of *S. canis* (Fig 2A). However, during logarithmic growth, we observe opacification activity caused by the bacterial sediment of the WT strain IMT49926 but not the *sof* deletion mutant, indicating that active ScSOF is covalently bound to the bacterial surface. Additionally, in SDS extracts of an overnight culture, we detect opacification activity in the wildtype strain but not in the deletion mutant, indicating that ScSOF is non-covalently associated with the surface in the stationary growth phase. Fig 2B and C show the zone of ß-haemolysis of IMT49926 and 49926Δ*sof* on a blood agar plate which is increased in the *sof* deletion mutant indicating that ScSOF could play a role in inhibition of ß-haemolysis.

**Fig 2:**
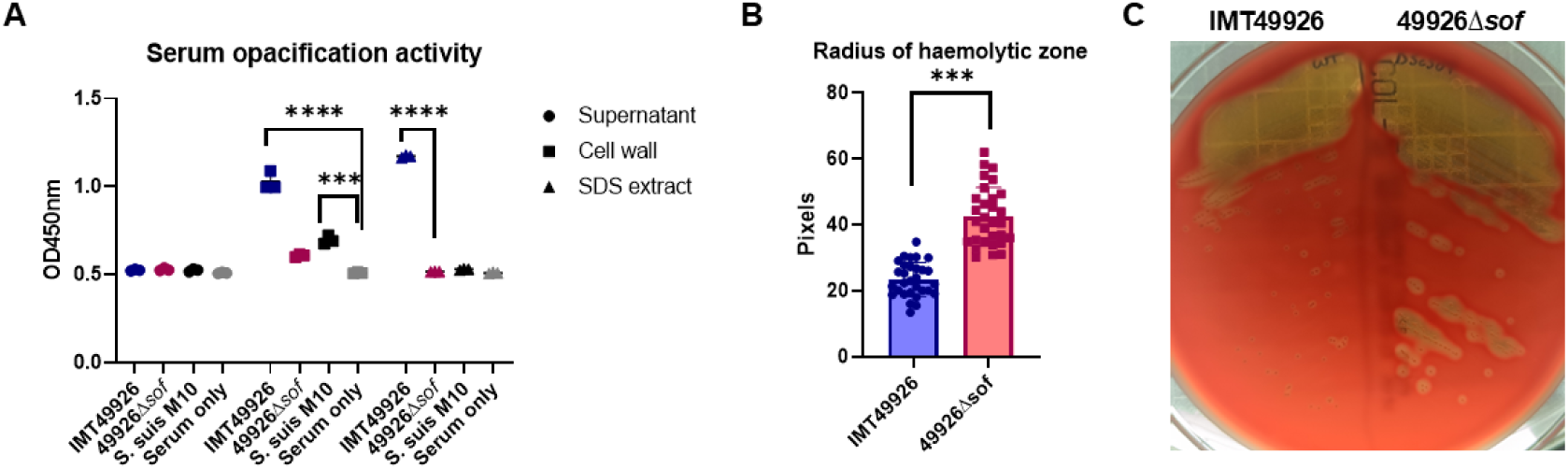
ScSOF opacifies serum and inhibits ß-haemolysis. A) Serum opacification activity of *S. canis* strain IMT49926 and 49926Δ*sof*, *S. suis* M10 and a negative serum control. Showing (left to right): overnight incubation with heat-inactivated bacterial supernatant, heat-inactivated bacterial sediment suspension from logarithmic growing bacteria and 1% SDS extracts of overnight cultures. B) Radius of the haemolytic zone measured with ImageJ based on pictures of blood agar plates of 3 biological replicates comparing IMT49926 and 49926Δ*sof* every time on the same plate. Statistical significance was calculated by Two-way ANOVA. *** indicates p < 0.001 and **** indicates p < 0.0001. C) Representative picture of a blood agar plate with colonies of *S. canis* IMT49926 (left) and 49926Δ*sof* (right).

### ScSOF is a major surface protein of *S. canis*

Protein sequence analyses of ScSOF discovered the presence of an N-terminal LPxTG-motif indicating surface expression of the SOF-protein. To visualize potential ScSOF-mediated differences in bacterial surface structures of *S. canis*, we used scanning electron microscopy to compare the WT strain IMT49926 with the mutant strain 49926Δ*sof*. *S. canis* IMT49926 exhibits a notable surface layer that appears rough and densely decorated (Fig 3A). The deletion mutant 49926Δ*sof* has a smoother, less textured surface, indicating loss of surface structures. To visualize the surface architecture of the wildtype and deletion mutant at a higher resolution, crosssectional images were acquired using a transmission electron microscopy as displayed in Fig 3B. Again, there is a notable difference in the thickness of the bacterial surface structures, displaying markedly decreased thickness for the mutant as confirmed by measurements of the surface material (mean surface thickness) shown in Fig 3C.

**Fig 3:**
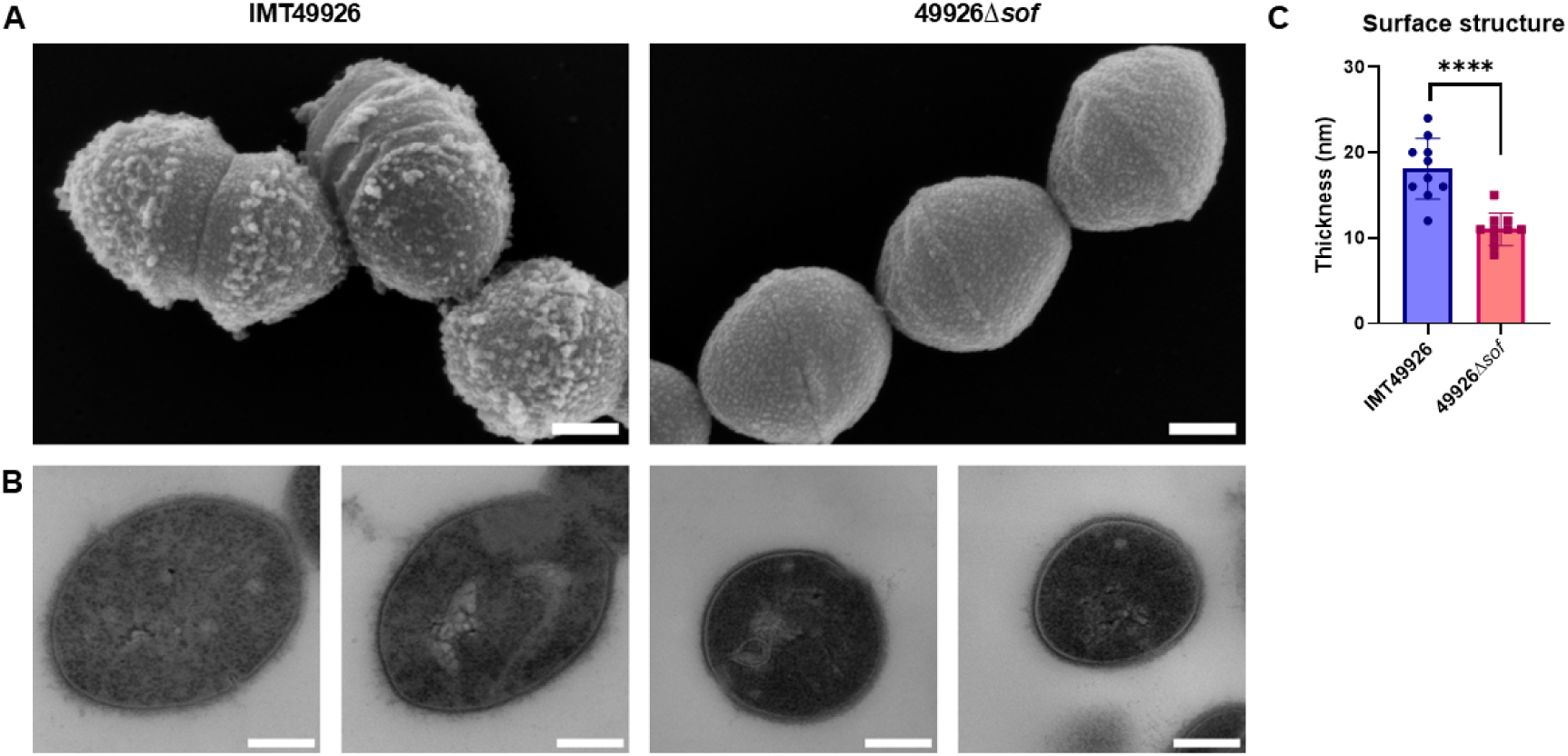
ScSOF is a major surface protein of *S. canis*. A) SEM visualization of IMT49926 in the left panel and of the 49926Δ*sof* mutant in the right panel. B) TEM images of IMT49926 and the 49926Δ*sof* mutant. Scale bars indicate 200 nm. C) Quantification of surface structure thickness of IMT49926 and 49926Δ*sof*. Statistical significance was calculated by Welch’s Students t-test. **** indicates p < 0.0001

### ScSOF is a fibronectin-binding protein

Domain homology comparisons identified three fibronectin-binding repeats in the protein sequence of ScSOF, pointing to a further important characteristic of ScSOF as a fibronectin-binding protein. To analyse the impact of ScSOF in fibronectin binding of *S. canis*, the wildtype strain and the isogenic ScSOF deletion mutant were used in two functional assays. First, immunofluorescence was used to visualize bacteria bound to a fibronectin-coated coverslip at different time points as shown in Fig 4A. After 2 hours, the *sof*-deletion mutant covers a substantially smaller fibronectin-coated area than the wildtype. However, after 3 hours, this difference is no longer visible. The quantification of fluorescence signals confirms the significant area reduction (p = 0.0218) of the fibronectin binding of the *sof*-deletion mutant after 2 hours (9.03 % ± 2.52 %) compared to the wildtype (18.9 % ± 3.95%) (Fig 4B). Second, binding of FITC-labelled recombinant fibronectin to wildtype and isogenic *sof*-deletion mutant was detected by FACS analyses. The geometric mean fluorescence intensity confirmed a slightly higher binding activity of wildtype bacteria to FITC-labelled fibronectin than of the *sof*-deletion mutant (Fig 4C).

**Fig 4:**
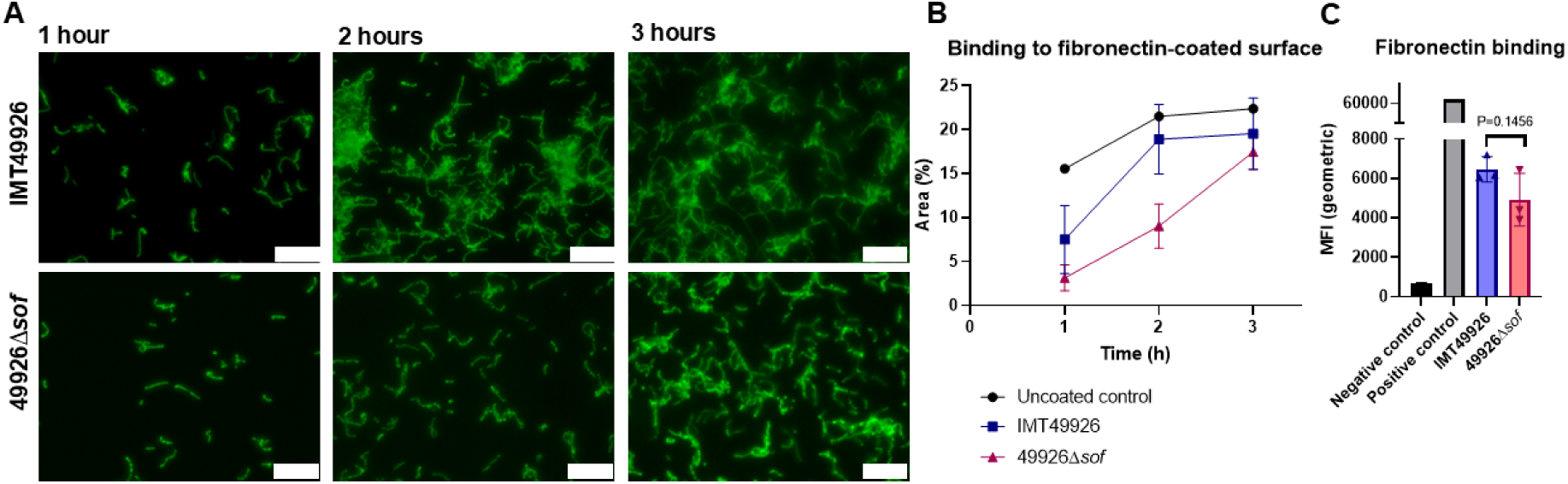
ScSOF plays a role in fibronectin-binding. A) *S. canis* IMT49926 and 49926Δ*sof* stained with specific antibodies (green) on a fibronectin-coated coverslip fixed at different timepoints. Images are acquired with a DMI6000B fluorescence microscope. Scale bars indicate 10 µm. B) Area covered with bacteria (determined by ImageJ analysis of images of three independent experiments. C) FACS analysis of the direct fibronectin-binding ability of IMT49926 and 49926Δ*sof* using FITC-conjugated fibronectin. Data represents geometric mean fluorescence intensity (MFI) ± SD of three independent experiments.

### Confirmation of the attenuated TraDIS mutant phenotype by the isogenic deletion mutant of the surface protein ScSOF

To confirm the attenuated phenotype of the transposon mutant, the isogenic deletion mutant was compared to the IMT49926 WT strain in the endothelial infection of HUVEC cells under flow conditions that was used to generate the TraDIS output pools. In the left panel of Fig 5A, uninfected HUVEC cells under flow conditions mimicking vascular shear stress are shown. The uninfected cells exhibit intact stress fibres and a tight cytoskeletal organization. In the middle panel, the wildtype infected cells show cytoskeletal remodelling including loss of stress fibres and formation of membrane protrusions and intracellular actin aggregates, with abundant bacterial association (yellow arrows). In the right panel, significantly reduced bacterial adherence and limited cytoskeletal perturbation is visualized after infection with the deletion mutant, comparable to the uninfected control. A quantification of the cell-associated bacteria also confirms a decrease in bacterial adherence to the endothelial cells (Fig 5B).

**Fig 5:**
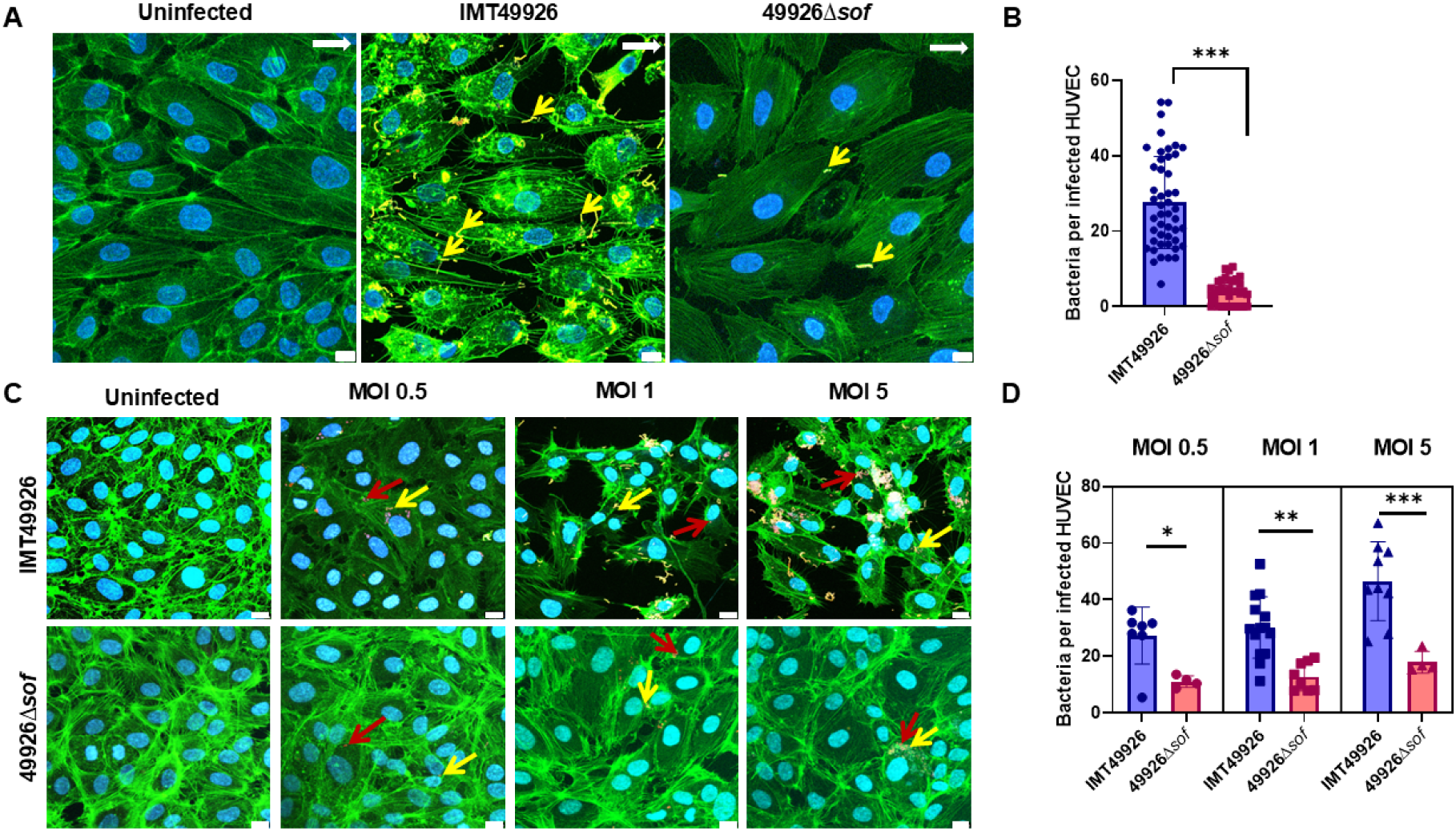
Deletion of the *sof* gene leads to a decrease in adherence and invasion of HUVECs. A) Representative images of HUVEC infection with IMT49926 and the 49926Δ*sof* mutant at MOI of 10 and shear stress of 6 dyn/cm² for up to 2.5 h (white arrows indicate direction of flow). Actin cytoskeleton was detected with green-fluorescent Alexa 488conjugated phalloidin. Nuclei were stained with DAPI (blue). Adherent streptococci appear in yellow and internalized bacteria in red. Images were acquired as z-stacks of 4.3 µm using CLSM (Leica, Sp8). Scale bars represent 10 µm. B) Quantification of cell-attached bacteria after infection under microfluidic conditions. C) HUVEC monolayers were infected with IMT49926 and the 49926Δ*sof* mutant at MOI 0.5, 1 and 5 under static conditions. Staining and imaging as above: actin cytoskeleton in green, nuclei in blue, adherent streptococci in yellow and internalized streptococci in red. Scale bars represent 10 µm. C) Quantification of bacterial adherence to HUVEC at different MOIs (0.5, 1, and 5) under static conditions. * Indicates p < 0.05, ** indicates p < 0.01, *** indicates p < 0.001.

To further analyse the impact of ScSOF in cell adherence, HUVECs were also infected under static conditions to exclude flow-induced host responses and the effect of shear stress on the bacterial adherence and cytoskeletal response. In Fig 5C, it is shown that with increasing MOI of the WT strain, there is an increase in cytoskeletal rearrangements, with disrupted actin filaments. There are multiple bacterial aggregates, with both adherent (yellow arrows) and internalized (red arrows) bacteria. Contrastingly, the HUVEC cells that were infected with the *sof* deletion mutant show no cytoskeletal differences compared to the uninfected control. The number of bacterial aggregates is severely decreased compared to WT infection. A quantification of the bacteria associated with the cells is shown for the different MOIs in Fig 5D showing a significant decrease in the number of bacteria that are adherent to or internalized into the HUVEC cells. A similar decrease in bacterial adherence and invasion was found upon infection of A549 lung epithelial cells (Fig S2), further demonstrating the importance of ScSOF in the pathogenicity of *S. canis*.

### ScSOF is involved in wound healing of endothelial cells

Septic bloodstream infections and infective endocarditis are linked to endothelial cell damage. Vascular shear stress regulates EC proliferation, migration, and barrier function. Using the Chamber-Separation cell Migration Assay (CSMA), HUVECs were exposed to increasing shear stress (3 to 10 dyn/cm² over 2 h), then infected with *S. canis* (Fig 6A-B). In controls, the cell-free gap closed progressively: 26.5% ± 9.1% at 6 h, 48% ± 12.7% at 24 h, and 99.98% ± 0.01% at 48 h. With WT infection, gap closure was severely impaired: only 8.3% ± 11.4% at 6 h and 0% at 24 h and 48 h, with the gap area even expanding due to cell loss. Infection with the *sof* deletion mutant allowed partial closure (24.2% ± 5.3% at 6 h; 49% ± 13.8% at 24 h), similarly to the control at 24 h (p = 0.99) as seen in Fig 6C. Statistical analysis confirmed significant differences between the uninfected control and the wildtype infected cells at all time points (p < 0.05), as well as between the wildtype infected cells and the mutant infected cells at 6 and 24 hours. However, at 48 hours, only the uninfected control reached complete gap closure, while both infected cells had an expanding gap area due to cell loss. Thus, SOF-positive *S. canis* strongly inhibits EC wound closure under physiological shear stress, while the mutant largely restores regeneration capacity at 24 hours, but not at 48 hours.

**Fig 6:**
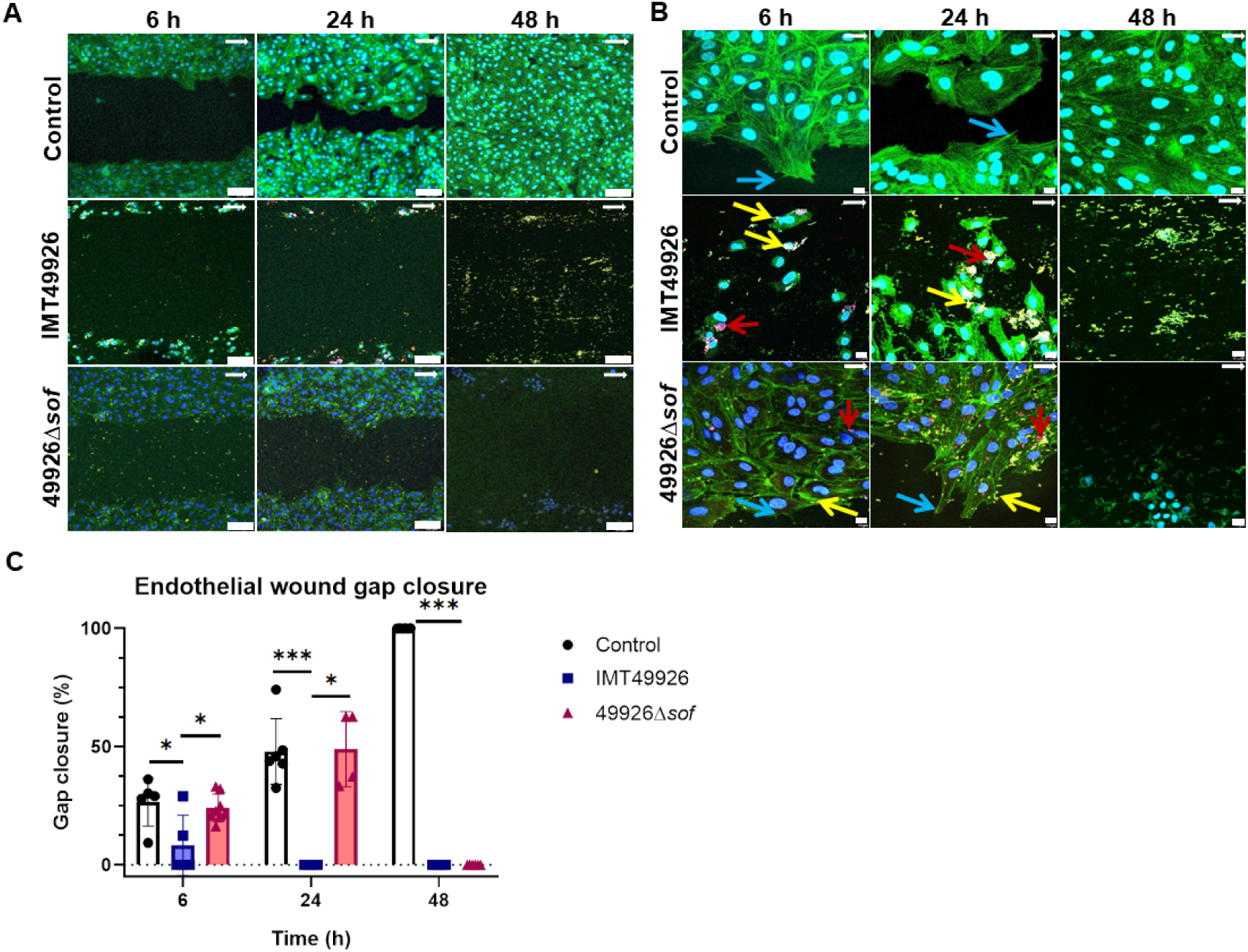
ScSOF plays a role in endothelial wound healing. A) Endothelial cell morphology was visualized by the detection of actin cytoskeleton with Alexa Fluor 488-conjugated phalloidin in combination with DAPI stain of nuclei at time points 6 h, 24 h and 48 h. Representative overlay images are shown for each time point of the CSMA without bacteria (ctrl), and in the presence of WT *S. canis* or 49926Δ*sof*. Images were generated using the CLSM (Leica Sp8) at 200-fold magnification and a zoom factor of 0.75. White arrows mark the direction of flow. Scale bar represents 100 μm. B) Representative microscopic images of CSMA results at 630-fold magnification. Blue arrows point to cell evaginations at the growing cell borders (ctrl). Yellow arrows point to adherent *S. canis*, and red arrows point to internalized bacteria. White arrows mark the direction of flow. Images generated using CLSM (Leica Sp8) at 630-fold magnification and a zoom factor of 0.75. Scale bar represents 10 μm. C) At indicated timepoints, progress of gap closure was calculated in percent in relation to the starting point, which is defined as 0% gap closure. Values are expressed as mean values with standard deviation. * Indicates p < 0.05, ** indicates p < 0.01, and *** indicates p < 0.001, according to one-way ANOVA and Tukey HSD post hoc tests. For the 24 h and 48 h samples homoscedasticity was not met (Levene p < 0.05); therefore, Welch’s ANOVA and Games-Howell post hoc testing were applied.

### *S. canis* inhibits cell proliferation and cell migration under physiological microfluidic conditions

Under microfluidic shear stress (10 dyn/cm²), HUVEC gap closure took up to 48 h. To determine how much of the gap closure was caused by cell proliferation and how much by cell migration, proliferating cells were labelled with EdU (Fig 7A). The proliferating cells (red nuclei) increased from 30.9 ± 21.6 at 6 h to 1101.7 ± 159 nuclei at 48 h, while migrating cells (blue nuclei) rose from 352.9 ± 184.4 to 541 ± 99.4 nuclei (Fig 7B-C). Proliferation dominated wound closure under flow conditions. Infection with WT *S. canis* significantly reduced proliferation (12.3 ± 8.3 at 6 h; 0.33 ± 0.47 at 24 h; none at 48 h) and migration (82 ± 25.7 at 6 h; 4.33 ± 5.4 at 24 h; none at 48 h) compared to the uninfected control. The *sof* deletion mutant showed proliferation and migration comparable to controls at 6 and 24 h, with significant differences from WT infection. These results demonstrate that *S. canis* impairs both endothelial cell proliferation and migration during wound healing under shear stress.

**Fig 7:**
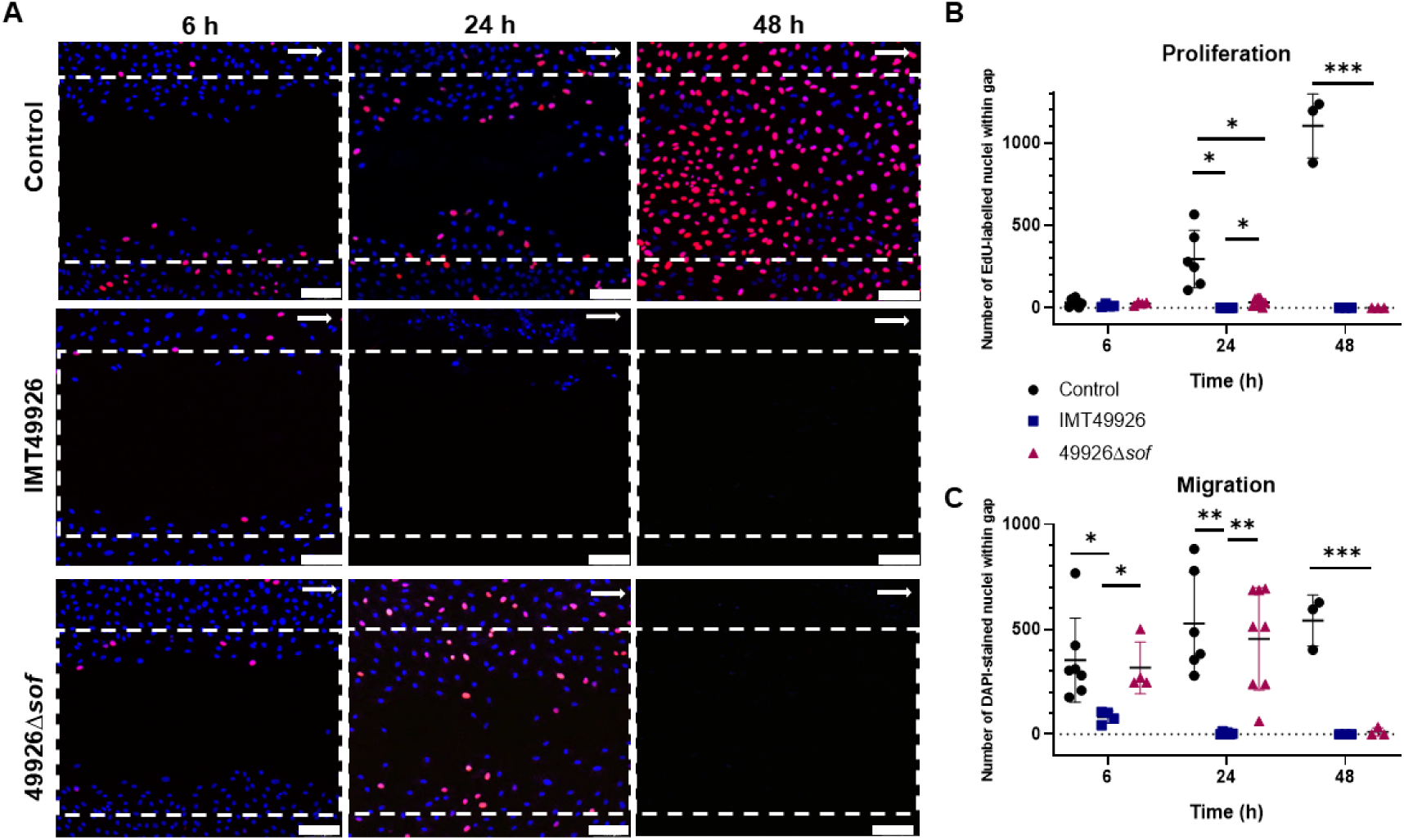
Differential microscopic evaluation of cell proliferation and migration with and without *S. canis* infection. A) CSMA was performed for indicated time periods without bacteria (control), and after infection with IMT49926 and 49926Δ*sof* at MOI 5 in the presence of EdU, which incorporates into proliferating HUVEC nuclei. In combination with 6-FAM-azide, nuclei of proliferating cells appear in light red, whereas migrating cells are counterstained in blue by DAPI incubation. In CSMA samples without bacteria, gap closure was completed after 48 h of cell cultivation. In the presence of bacteria, the cell-free gaps were not closed after 48 h of cell cultivation. Representative microscopic fluorescence images are shown for each sample. Images were generated using the CLSM (Leica Sp8) at 200-fold magnification and a zoom factor of 0.75. Scale bar represents 100 μm. Proportion of proliferating HUVECs (B) and migrating cells (C) was quantified by counting the red and blue fluorescing cell nuclei in the defined gap area at indicated time points (6 h, 24 h and 48 h. Experiments were performed in three independent assays, each in triplicate, and a minimum of four different fields of view were randomly chosen for quantification of cell nuclei. Data are expressed as mean value including standard deviation. * Indicates p < 0.05, ** indicates p < 0.01, and *** indicates p < 0.001, according to one-way ANOVA and Tukey HSD post hoc tests. For the 24 h samples *homoscedasticity was not met (Levene p < 0.05); therefore, Welch’s ANOVA and Games-Howell post hoc testing were applied*.

## Supplementary figures

**S1: Insufficient separation of non-adherent and adherent transposon mutant after first infection round.** Volcano plot of a comparative analysis of the generated output pool (nonadherent bacteria after first selection round) and the input library showing the logFC of all genes after the first round of infection compared to the input pool (control).

**S2: A549 epithelial cells after infection with IMT49926 and 49926Δ*sof* at MOI 5.** Staining done as in the HUVEC infection experiment, but phalloidin staining is visualized in white. Adherent bacteria are shown in yellow and internalized bacteria in red as before. Scale bar represents 10 µm. Imaged with a DMI6000B fluorescence microscope at 630x magnification. A quantification of the total CFU after infection is shown on the right. *** indicates p < 0.001.

## Discussion

Despite increasing recognition of *S. canis* as a causative agent of infective endocarditis (IE) in both dogs and humans, the molecular mechanisms underlying its adherence to the host endothelium remain uncharacterized. While factors such as the M protein SCM have been implicated in immune evasion and early colonization, other specific adherence factors and their roles in endothelial interactions remain poorly understood [44]. Given the importance of fibronectin-binding proteins in the virulence of related streptococcal pathogens, it is plausible that similar mechanisms are at play in *S. canis*, but direct evidence has been lacking [19, 39].

We employed a high-throughput TraDIS approach in combination with a physiologically relevant *in vitro* endothelial infection model that incorporates shear stress, thus mimicking the dynamic conditions of the bloodstream. This strategy allowed us to identify key genes involved in endothelial adherence.

The transposon mutant library provided sufficient genome coverage and highresolution mapping of gene function. The comparison between input and nonadherent output pools revealed significant shifts in mutant abundance, indicating efficient selection pressure and successful enrichment for adherence-associated genes. We identified multiple genes potentially involved in adherence and invasion of endothelial cells. Only three LPXTG-motif containing surface proteins passed our applied threshold (logFC > 2, q < 0.01), suggesting they are critical for adherence. Among these, a fibronectin-binding protein with high homology to known serum opacity factors (SOFs) emerged as a promising candidate for further functional validation. ScSOF was selected due to its predicted fibronectin-binding capacity and homology to virulence-associated SOFs in other streptococcal species.

Deletion of *sof* confirmed the role of ScSOF in mediating endothelial cell adherence and invasion. Under flow conditions that mimic vascular shear stress, wildtype IMT49926 induced substantial cytoskeletal rearrangement and bacterial adherence, whereas the *sof* deletion mutant showed significantly reduced attachment and minimal host cytoskeletal disruption. This phenotype was consistently observed under static conditions and at different MOIs, confirming the importance of ScSOF for endothelial cell adherence, independent of flow-induced responses.

The signature characteristic of SOF proteins is their ability to opacify serum. Our data show that cell-associated ScSOF contributes to serum opacification, with both covalent and non-covalent associations depending on the bacterial growth phase. Additionally, the *sof* deletion mutant displayed enhanced β-haemolysis on blood agar, implicating a role for ScSOF in the modulation of haemolytic activity. This suggests that ScSOF has similar properties as SOF of *S. pyogenes*, which is also described as inhibiting β-haemolysis [41].

Electron microscopy highlighted the role of ScSOF as a major surface-associated factor. SEM and TEM imaging demonstrated a significant reduction in the thickness and complexity of surface structures in the *sof* deletion mutant compared to the wildtype, suggesting ScSOF contributes either directly to the extracellular matrix or indirectly by recruiting other surface-associated materials.

Binding assays validated the function of ScSOF as a fibronectin-binding protein. Flow cytometry with FITC-labelled fibronectin and immunofluorescence microscopy showed a reduction in fibronectin binding by the 49926Δ*sof* strain, particularly at earlier timepoints, reinforcing the importance of ScSOF in initiating host interactions. Although this difference diminishes over time, likely due to compensatory mechanisms by other fibronectin-binding surface proteins or nonspecific binding, early adhesion events are critical in pathogenesis, particularly for infective endocarditis (IE), where initial endothelial attachment is a prerequisite for disease progression [45]. Other binding characteristics of ScSOF could be tested in further studies, as SOF of *S. pyogenes* has for example been shown to bind to fibulin-1, another extracellular host matrix protein [40].

Consistent with reduced cell attachment, CSMA analyses showed that the *sof* deletion mutant had significantly better cell layer generation compared to the WT. In the absence of infection, HUVEC monolayers fully closed the cell-free gap within 48 h under microfluidic conditions. Infection with WT severely impaired both proliferation and migration, causing wound expansion rather than closure. In contrast, the *sof* deletion mutant allowed significantly increased regeneration compared to the WT after 24 hours, but both infected conditions inhibited wound closure after 48 hours.

The reduced adherence of the mutant likely contributes to its diminished impact on endothelial repair. Overall, these results demonstrate that SOF expression in *S. canis* is a key factor in disrupting endothelial wound healing under flow conditions, making it an important bacterial factor in the development of infective endocarditis.

We further showed that the *sof* deletion mutant had a decreased adherence to epithelial cells using A549 lung epithelial cells. This is in line with results from the SOF protein of *S. pyogenes*, which was shown to play a role in adherence to different epithelial cell lines, mainly by adherence to the host target matrix component fibronectin [42]. It was also shown for *S. pyogenes* that next to the decrease in adherence to epithelial cells, elimination of SOF led to a decrease of virulence in a murine model of necrotizing skin infection [46]. Because the ScSOF has a high similarity to the SOF of *S. pyogenes*, it is likely that ScSOF plays a similar role in *S. canis*, where a decrease in virulence could be expected upon deletion of the *sof* gene.

Overall, our findings identified ScSOF as a multifunctional surface protein of *S. canis* that plays a role in IE pathophysiology. It facilitates adhesion to endothelial cells, contributes to surface architecture, binds fibronectin, opacifies serum, and inhibits βhaemolytic activity. These properties make ScSOF a compelling candidate for further studies into vaccine development or therapeutic targeting.

Future work should explore the precise molecular interactions between ScSOF and fibronectin and other host matrix proteins such as fibulin, the structural biology of ScSOF itself, and its regulation under *in vivo* conditions. Additionally, given its similarity to SOFs in *S. pyogenes* and *S. dysgalactiae*, comparative functional analyses may reveal conserved virulence strategies across pyogenic streptococci. The two other genes that were identified in the TraDIS analysis, an IgG-binding protein G homologue and the M-like protein SCM type II, are of interest for further research too. Streptococcal M proteins are key virulence factors and often found to play a role in the interaction with host cells and host matrix proteins [14–16, 47]. The IgG-binding protein G homologue could potentially bind IgG and albumin, properties that can promote formation of vegetations by facilitating adhesion to fibrin-platelet aggregates on damaged heart valves [48–50].

## Acknowledgements

The authors thank Nina Janze and Ina Brentrop for the excellent technical assistance. Furthermore, we thank the Sequencing Core Facility of the Genome Competence Centre (MF1), Robert Koch Institute for providing excellent support in TraDIS sequencing.

This work was supported by the German Research Foundation (DFG) (grant FU 1027/3-1 to MF and FU 1027/5-1 to SB, MF).

## CRediT authorship contribution statement

Conceptualization, M.F., S.B.; methodology, M.F., S.B., A.K. and M.K.; software, E.A., S.A.W.,M.K.; validation, G.R., D.S., S.H., A.K. and M.K.; formal analysis, A.K., M.K.; investigation, G.R., D.S., S.H., E.A., M.M., A.K. and M.K.; resources, M.M., I.E., S.B. and M.F.; data curation, E.A., S.A.W., A.K. and M.K.; writing—original draft preparation, A.K. and M.K.; writing—review and editing, all; visualization, A.K. and M.K.; supervision, S.B. and M.F.; project administration, S.B. and M.F.; funding acquisition, S.B. and M.F. All authors have read and agreed to the published version of the manuscript.

## Data availability statement

Sequencing data have been deposited in the NCBI BioProject repository and are publicly available under BioProject accession: PRJNA945807, BioSample accessions SAMN53665527-SAMN53665533. Other relevant data is provided in the supporting information file.

## Reference List

1. Lysková, P., et al., Prevalence and Characteristics of Streptococcus canis Strains Isolated from Dogs and Cats. Acta Veterinaria Brno, 2007. 76(4): p. 619–625.

2. Lamm, C., et al., Streptococcal infection in dogs: a retrospective study of 393 cases. Veterinary pathology, 2010. 47(3): p. 387–395.

3. Sykes, J.E., et al., Evaluation of the relationship between causative organisms and clinical characteristics of infective endocarditis in dogs: 71 cases (1992– 2005). Journal of the American Veterinary Medical Association, 2006. 228(11): p. 1723–1734.

4. Amsallem, M., et al., First reported human case of native mitral infective endocarditis caused by Streptococcus canis. Canadian Journal of Cardiology, 2014. 30(11): p. 1462. e1-1462. e2.

5. Lacave, G., et al., Endocarditis caused by Streptococcus canis: an emerging zoonosis? Infection, 2016. 44: p. 111–114.

6. Mališová, B., et al., Human native endocarditis caused by Streptococcus canis—a case report. Apmis, 2019. 127(1): p. 41–44.

7. Wang, M.S., et al., Streptococcus canis Native Aortic Valve Endocarditis Linked to Cat Exposure: A Case Report and Review. Journal of Community Hospital Internal Medicine Perspectives, 2024. 14(2): p. 91.

8. MacDonald, K., Infective endocarditis in dogs: diagnosis and therapy. Veterinary Clinics: Small Animal Practice, 2010. 40(4): p. 665–684.

9. Katsburg, M., et al., Limiting Factors in Treatment Success of Biofilm-Forming Streptococci in the Case of Canine Infective Endocarditis Caused by Streptococcus canis. Veterinary Sciences, 2023. 10(5): p. 314.

10. Reagan, K.L., et al., Outcome and prognostic factors in infective endocarditis in dogs: 113 cases (2005-2020). Journal of Veterinary Internal Medicine, 2022. 36(2): p. 429–440.

11. Moser, C., et al., Biofilms and host response–helpful or harmful. Apmis, 2017. 125(4): p. 320–338.

12. Witten, J.C., et al., Basic Science Aspects of the Pathogenesis of Infective Endocarditis. Infective Endocarditis, 2024.

13. Lerche, C.J., et al., Anti-biofilm approach in infective endocarditis Exposes new treatment strategies for improved outcome. Frontiers in Cell and Developmental Biology, 2021. 9: p. 643335.

14. Bergmann, S., et al., SCM, the M protein of Streptococcus canis binds immunoglobulin G. Frontiers in Cellular and Infection Microbiology, 2017. 7: p. 80.

15. Fulde, M., et al., SCM, a novel M-like protein from Streptococcus canis, binds (mini)-plasminogen with high affinity and facilitates bacterial transmigration. Biochemical Journal, 2011. 434(3): p. 523–535.

16. Lapschies, A.-M., et al., The type-2 Streptococcus canis M protein SCM-2 binds fibrinogen and facilitates antiphagocytic properties. Frontiers in Microbiology, 2023. 14: p. 1228472.

17. Cornax, I., et al., Novel Models of Streptococcus canis Colonization and Disease Reveal Modest Contributions of M-Like (SCM) Protein. Microorganisms 2021, Vol. 9, Page 183, 2021-01-16. 9(1).

18. Widmer, E., et al., New concepts in the pathophysiology of infective endocarditis. Current Infectious Disease Reports 2006 8:4, 2006/08. 8(4).

19. Que, Y.-A., et al., Fibrinogen and fibronectin binding cooperate for valve infection and invasion in Staphylococcus aureus experimental endocarditis. Journal of Experimental Medicine, 2005/05/16. 201(10).

20. Jung, C.-J., et al., Streptococcus mutans autolysin AtlA is a fibronectin-binding protein and contributes to bacterial survival in the bloodstream and virulence for infective endocarditis. Molecular Microbiology, 2009/11/01. 74(4).

21. Oppegaard, O., et al., Clinical and molecular characteristics of infective βhemolytic streptococcal endocarditis. Diagnostic microbiology and infectious disease, 2017. 89(2): p. 135–142.

22. Bedran, T.B.L., et al., Fibrinogen-Induced Streptococcus mutans Biofilm Formation and Adherence to Endothelial Cells. BioMed Research International, 2013/01/01. 2013(1).

23. GC, L., et al., Simultaneous assay of every Salmonella Typhi gene using one million transposon mutants - PubMed. Genome research, 2009 Dec. 19(12).

24. Charbonneau, A.R.L., et al., Identification of genes required for the fitness of Streptococcus equi subsp. equi in whole equine blood and hydrogen peroxide. Microbial Genomics, 2020/03/31. 6(4).

25. Barquist, L., et al., The TraDIS toolkit: sequencing and analysis for dense transposon mutant libraries. Bioinformatics, 2016/04/01. 32(7).

26. Charbonneau, A.R.L., et al., Defining the ABC of gene essentiality in streptococci. BMC Genomics 2017 18:1, 2017–05-31. 18(1).

27. Moule, M.G., et al., Characterization of New Virulence Factors Involved in the Intracellular Growth and Survival of Burkholderia pseudomallei. Infection and Immunity, 2015-12-28. 84(3).

28. Weerdenburg, E.M., et al., Genome-Wide Transposon Mutagenesis Indicates that Mycobacterium marinum Customizes Its Virulence Mechanisms for Survival and Replication in Different Hosts. Infection and Immunity, 2015-2-17. 83(5).

29. Jagau, H., et al., Frontiers | Von Willebrand Factor Mediates Pneumococcal Aggregation and Adhesion in Blood Flow. Frontiers in Microbiology, 2019/03/26. 10.

30. Baums, C.G., et al., Identification of a Novel Virulence Determinant with Serum Opacification Activity in Streptococcus suis. Infection and Immunity, 2006-November. 74(11).

31. Kopenhagen, A., et al., Frontiers | Streptococcus pneumoniae Affects Endothelial Cell Migration in Microfluidic Circulation. Frontiers in Microbiology, 2022/03/25. 13.

32. Bolger, A.M., M. Lohse, and B. Usadel, Trimmomatic: a flexible trimmer for Illumina sequence data. Bioinformatics, 2014/08/01. 30(15).

33. Rutherford, K., et al., Artemis: sequence visualization and annotation. Bioinformatics, 2000/10/01. 16(10).

34. Carver, T., et al., DNAPlotter: circular and linear interactive genome visualization. Bioinformatics, 2009/01/01. 25(1).

35. Cantalapiedra, C.P., et al., eggNOG-mapper v2: Functional Annotation, Orthology Assignments, and Domain Prediction at the Metagenomic Scale. Molecular Biology and Evolution, 2021/12/09. 38(12).

36. Deschner, F., et al., Natural products chlorotonils exert a complex antibacterial mechanism and address multiple targets. Cell Chemical Biology, 2025/04/17. 32(4).

37. Courtney, H.S. and H.J. Pownall, The Structure and Function of Serum Opacity Factor: A Unique Streptococcal Virulence Determinant That Targets High-Density Lipoproteins. BioMed Research International, 2010/01/01. 2010(1).

38. Fischetti, V.A., Surface Proteins on Gram-Positive Bacteria. Gram-Positive Pathogens, 2014.

39. Courtney, H.S., et al., Serum opacity factor is a major fibronectin-binding protein and a virulence determinant of M type 2 Streptococcus pyogenes. Molecular microbiology, 1999. 32(1): p. 89–98.

40. Courtney, H.S., et al., Serum Opacity Factor Is a Streptococcal Receptor for the Extracellular Matrix Protein Fibulin-1 *. Journal of Biological Chemistry, 2009/05/08. 284(19).

41. Zhu, L., R.J. Olsen, and J.M. Musser, Opacification Domain of Serum Opacity Factor Inhibits Beta-Hemolysis and Contributes to Virulence of Streptococcus pyogenes. mSphere, 2017-4-19. 2(2).

42. Oehmcke, S., A. Podbielski, and B. Kreikemeyer, Function of the FibronectinBinding Serum Opacity Factor of Streptococcus pyogenes in Adherence to Epithelial Cells. Infection and Immunity, 2004-7. 72(7).

43. Katerov, V., et al., Streptococcal Opacity Factor: A Family of Bifunctional Proteins with Lipoproteinase and Fibronectin-Binding Activities. Current Microbiology 2000 40:3, 2000/03. 40(3).

44. Pagnossin, D., et al., Streptococcus canis, the underdog of the genus. Veterinary Microbiology, 2022/10/01. 273.

45. Chorianopoulos, E., et al., The role of endothelial cell biology in endocarditis. Cell and Tissue Research 2008 335:1, 2008–11-18. 335(1).

46. Timmer, A.M., et al., Serum opacity factor promotes group A streptococcal epithelial cell invasion and virulence. Molecular Microbiology, 2006/10/01. 62(1).

47. Oehmcke, S., et al., Streptococcal M proteins and their role as virulence determinants. Clinica Chimica Acta, 2010/09/06. 411(17-18).

48. Isobel Ford, C.W.I.D., The role of platelets in infective endocarditis. Platelets, 1997-1-1. 8(5).

49. Talay, S.R., M.P. Grammel, and G.S. Chhatwal, Structure of a group C streptococcal protein that binds to fibrinogen, albumin and immunoglobulin G via overlapping modules. Biochemical Journal, 1996/04/15. 315(2).

50. Nygren, P.-Å., et al., Analysis and use of the serum albumin binding domains of streptococcal protein G. Journal of Molecular Recognition, 1988/04/01. 1(2).

